# Structural plasticity and enhanced fear extinction following psilocybin in chronically stressed mice

**DOI:** 10.64898/2026.04.21.720014

**Authors:** Cory A. Knox, Samuel C. Woodburn, Amelia D. Gilbert, Jason M. Schlotzhauer, Alex C. Kwan

## Abstract

The classic psychedelic psilocybin elicits long-lasting neural plasticity and behavioral effects, but prior studies largely examined stress-naive animals. Using longitudinal imaging, we show that psilocybin increases dendritic spine density in frontal cortical neurons and facilitates fear extinction after chronic restraint stress, demonstrating psilocybin’s effects in a translationally relevant mouse model.

## Introduction

Structural neural plasticity likely underpins the effects of rapid-acting antidepressant interventions ^1^. Synaptic dysfunction is observed in the prefrontal cortex of individuals with depression ^2, 3^. Dendritic atrophy and dendritic spine loss are present in the medial frontal cortex of rodent stress models ^4, 5^. By contrast, the rapid-acting antidepressant ketamine increases the dendritic spine numbers in rat medial frontal cortex ^6–8^. Recently, classic psychedelics and related serotonin 2A (5-HT_2A_) receptor agonists have also been shown to promote structural plasticity ^9, 10^. Notably, one dose of psilocybin rapidly induces dendritic spine formation in layer 5 pyramidal cells of the mouse medial frontal cortex ^11–15^. The effect is long-lasting, detectable within 24 hr and persisting for at least 1 month in the mouse brain ^13^. Structural plasticity appears to be a shared feature of psychedelics, as the effect generalizes to other compounds including 5-MeO-DMT ^16^, DOI ^17^, 25CN-NBOH ^18^, as well as various non-hallucinogenic analogs or serotonergic manipulations ^11, 19–23^.

However, a key limitation of most prior studies for psilocybin is that drug-evoked structural plasticity was assessed in stress-naïve, wild type animals. In preclinical research, chronic stress models are used to induce aspects of psychiatric pathophysiology, because repeated stress exposure is a major risk factor for many disorders including depression. It is possible for the effects of pharmacological interventions to differ between unstressed and stressed animals. Indeed, subanesthetic dose of ketamine has been reported to reduce immobility in forced swim test for male mice subjected to unpredictable chronic stress, while exacerbating the behavioral measure in unstressed animals ^24^. Divergent effects have been observed in humans, where ketamine produced opposite changes in depressive symptoms in individuals with major depressive order compared to healthy controls ^25^. These findings underscore the importance of extending studies of psilocybin-induced structural plasticity to chronic stress models.

## Results

Stress can impair the extinction of cued fear ^26^, but psilocybin has the opposite effect of facilitating fear extinction ^27, 28^. Here, to investigate the potential interaction between stress and psilocybin, C57BL/6J mice were subjected to restraint every day for two weeks (day −14 to −1). Control mice were handled but not restrained. A total of 24 animals subsequently went through fear conditioning on day 0, and received psilocybin (1 mg/kg, i.p.) or saline prior to fear extinction on day 2. This timing in which psilocybin is psychoactive during fear extinction is key for the long-term behavioral effects, as our previous study has shown by varying the timing of administration ^27^. Afterwards, the cohort was divided with 16 mice sacrificed immediately for immunohistochemistry to assess activity-dependent transcription in the medial frontal cortex and 8 mice tested for fear renewal testing on day 10 (**Fig. 1A**). Weight was recorded on day −15 and day 0, which showed a significant difference between the controls and stressed mice (*P*=1.4×10^-22^, two-tailed t-test; **Fig. 1B**). We assessed cells labeled by neuronal excitability marker phosphorylated cAMP response element-binding protein (pCREB) in three regions: dorsal anterior cingulate (ACAd), prelimbic (PL), and infralimbic (ILA) (**Fig. 1C**). We found that pCREB+ cell density differed across region (*P*<0.0001, generalized linear mixed effects model), but did not detect effects of stress (*P*=0.16), drug (*P*=0.85), or stress-drug interaction (*P*=0.31). For control and vehicle, control and psilocybin, stress and vehicle, and stress and psilocybin groups, the pCREB+ cell densities were 2586±80, 2731±369, 2449±90, 2229±104 mm^-2^ in ACAd; 2549±75, 2694±336, 2469±72, 2290±87 mm^-2^ in PL; and 2778±117, 2948±370, 2770 ±100, 2504±106 mm^-2^ in ILA, respectively (**Fig 1D-F**).

**Fig. 1.**
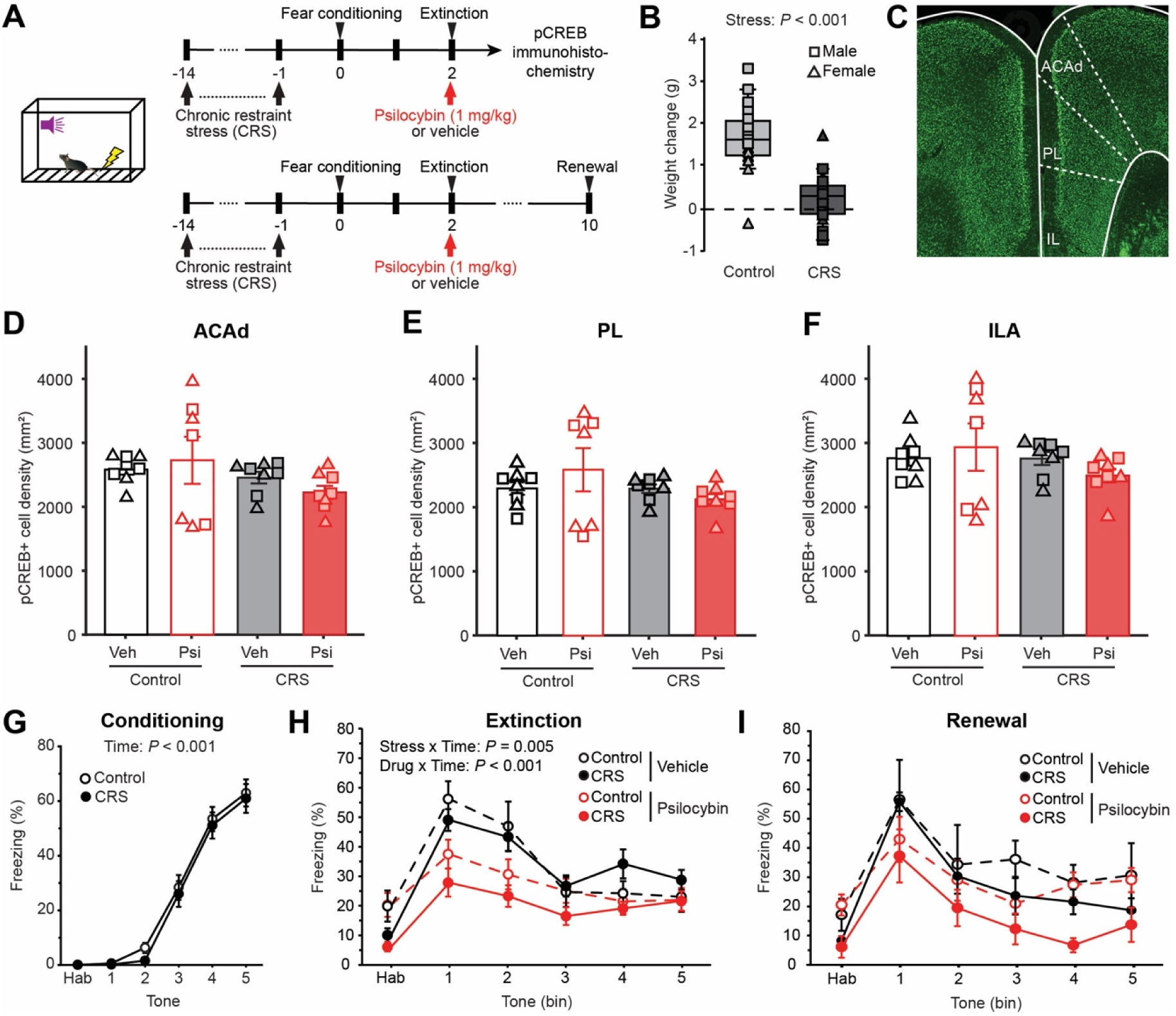
Enhanced fear extinction following psilocybin in chronically stressed mice. **(A)** Experimental timeline. **(B)** Change in weight from day −15 to day 0 for animals in the control unstressed and chronic restraint stress (CRS) groups. Square, male. Triangle, female. **(C)** Fluorescence image of a coronal section with pCREB-positive cells. **(D)** Density of pCREB-positive cells in the dorsal anterior cingulate area (ACAd). Square, male. Triangle, female. Bar, mean. Error bars, ±SEM. **(E)** Similar to (D) for prelimbic area (PL). **(F)** Similar to (D) for infralimbic area (ILA). **(G)** Fraction of time spent freezing during different CS presentations during fear conditioning on day 0. Line, mean. Error bars, ±SEM. **(H)** Fraction of time spent freezing during habituation (Hab) and different binned CS presentations during extinction on day 2. Line, mean. Error bars, ±SEM. **(I)** Similar to (E) for fear renewal on day 10.

In terms of freezing behavior, we did not detect any difference between stressed mice and unstressed controls in their ability to develop fearful associations (**Fig. 1G**). During extinction, mice given psilocybin exhibited less freezing than vehicle-treated animals (drug × time interaction: *P*=0.041, drug × stress interaction: *P*=0.02, generalized linear mixed effects model; **Fig. 1H**). Post hoc analysis showed that psilocybin’s effects were significant for both controls (*P*=0.0003, sequential Bonferroni analysis) and stressed mice (*P*=0.0001). Fear renewal on day 10 showed trends that are similar to extinction on day 2, but the statistical comparisons did not yield significant differences, likely owing to the smaller sample size (**Fig. 1I**). Together, these results demonstrate that psilocybin enhances fear extinction in chronically stressed mice to a similar degree as in stress-naïve mice.

To measure structural plasticity, we implanted a cranial window for two-photon imaging of GFP-expressing apical dendrites in the medial frontal cortex of *Thy1^GFP^* line M mice (**Fig. 2A**). The same dendrites were tracked across 7 imaging sessions on day −15, −3, −1, 1, 3, 5, and 14, while chronic restraint stress was applied from day −14 to day −1, and psilocybin (1 mg/kg, i.p.) or vehicle was administered on day 0 (**Fig. 2B, C**). In total, we counted 56, 68, and 57 dendrites including 606, 836, and 484 dendritic spines in the three experimental groups: stress/psilocybin, stress/vehicle, and control/vehicle, respectively. We plotted the fold-change in spine density, using pre-treatment value (average of values on day −3 and day −1) as the baseline (**Fig. 2D**). For statistical analysis, we used a mixed-effects model that accounts for the fact that multiple dendrites were measured from the same mouse (see **Methods**). Notably, there was a significant interaction effect for group and time (*P*=2×10^−11^, mixed effects model), indicating that spine density varied across days differentially for the three experimental groups.

**Fig. 2.**
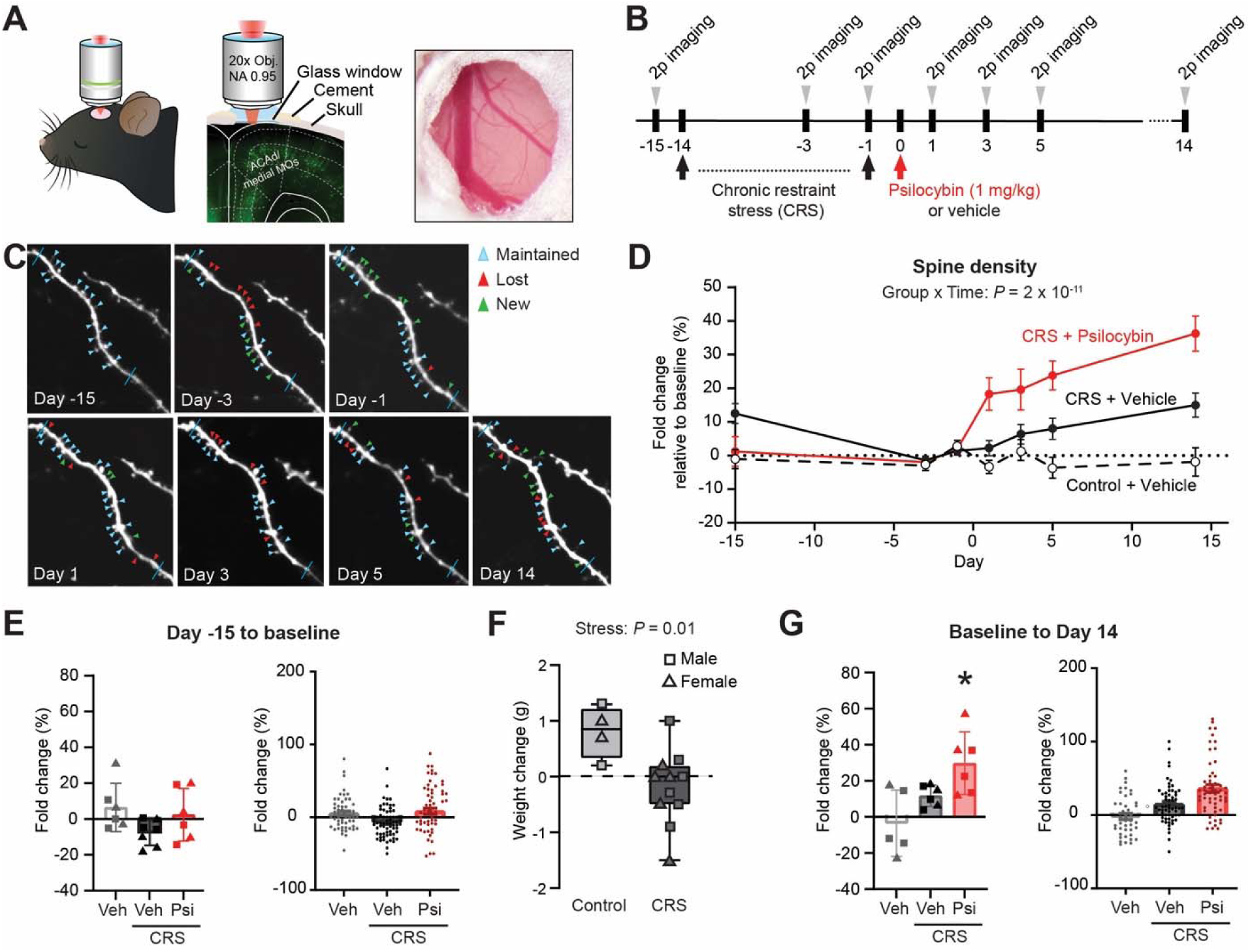
Structural plasticity following psilocybin in chronically stressed mice. **(A)** Imaging setup. **(B)** Experimental timeline. **(C)** Two-photon image of a field of view, tracking the same apical tuft dendrites in *Thy1^GFP^* mice across imaging sessions. **(D)** Dendritic spine density as a function of time. Spine density was expressed as fold-change from baseline (average of values on on day −3 and day −1) for each dendritic segment. Line, mean. Error bars, ±SEM. **(E)** Fold-change in dendritic spine density from day −15 to baseline. Fold-change is calculated with value on day −15 as the denominator. Left, per mouse. Right, per dendrite. Line, mean. Error bars, ±SEM. **(F)** Change in weight from day −15 to day 0 for *Thy1^GFP^* mice in the control unstressed and CRS groups. Square, male. Triangle, female. **(G)** Similar to (K), from baseline to day 14. Fold-change is calculated with value at baseline (average of values on day −3 and day −1) as the denominator. *, *P* < 0.05; post hoc test via estimated marginal means with Tukey correction.

We were particularly interested in the effects of stress and psilocybin. To assess the impact of stress, we compared spine density between day −15 and baseline, plotting either on a per-animal (left panel, **Fig. 2E**) or per-dendrite basis (right panel, **Fig. 2E**). Statistical analysis revealed no main effect of group (*P*=0.2, mixed effects model), indicating no overt impact of chronic restraint stress on dendritic spine density. We were concerned that the chronic restraint stress might be ineffective for the *Thy1^GFP^* mice, yet this was unlikely because there was a clear difference in weight change between control and stressed mice (*P*=0.016, unpaired t-test; **Fig. 2F**). To assess the impact of psilocybin, we compared spine density between baseline and day 14 (**Fig. 2G**). This analysis revealed a significant main effect for group (*P*=0.012, mixed effects model). Specifically, post hoc analysis revealed that there was a significant increase for the stress and psilocybin group, relative to the control and vehicle and stress and vehicle groups (*P*=0.0023, estimated marginal means with FDR correction). Collectively, the results show that a single dose of psilocybin increases dendritic spine density in chronically stressed mice.

To determine whether psilocybin-induced increase in spine density reflected enhanced formation, reduced elimination, or both, we analyzed the turnover of dendritic spines between consecutive imaging sessions. Formation and elimination rates were visualized as the difference from the pre-treatment rates (from day −3 to day −1). The spine formation rate was higher in the CRS + Psilocybin group than in the CRS + Vehicle group (*P* = 0.012 on day 1 and *P* = 0.031 on day 5, estimated marginal means with FDR correction following a mixed-effects model; **Fig. 3A**). By contrast, spine elimination rates did not differ among the three groups (**Fig. 3B**). Thus, the increase in spine density following psilocybin in chronically stressed mice arose mainly from enhanced spine formation rather than reduced spine elimination.

**Figure 3:**
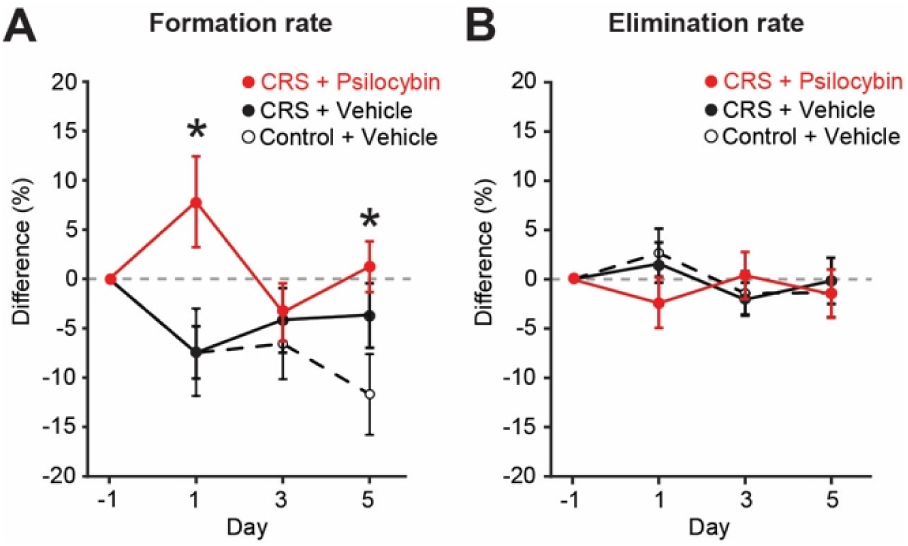
Turnover dynamics of dendritic spines following psilocybin in chronically stressed mice. **(A)** The formation rates of dendritic spines, plotted as difference from baseline (formation rate between sessions on day −3 and day −1). Line, mean. Error bars, ± SEM. **(B)** Similar to (A) for elimination rates. Line, mean. Error bars, ±SEM.

## Discussion

Prior studies reported varied effects of chronic stress, such as enhanced fear learning and impaired fear extinction ^29^, or spared learning and extinction but impaired extinction recall ^30^. Moreover, in prior studies of stressed animals using *in vivo* two-photon imaging, spine elimination was observed following chronic corticosterone exposure ^31^ or unpredictable mild stress ^20^. Differences among prior studies and with our results could arise due to the type and duration of stressor, biological variables such as sex, or the cortical region examined for structural plasticity. Of note, the time courses for the CRS + vehicle and control + vehicle groups in **Fig. 2D** suggest a possible rebound increase in spine density following the end of the chronic restraint period. Therefore, although we did not detect a direct effect of chronic stress on spine density, the subsequent increase in the stressed group relative to controls suggests that structural remodeling may occur during the recovery phase.

Excitatory pyramidal neurons comprise multiple genetically distinct classes, including pyramidal tract (PT), intratelencephalic (IT), and corticothalamic (CT) neurons ^32, 33^. These classes differ in their physiology, connectivity, and contributions to behavior ^34–36^. Recent work has shown that psilocybin induces structural remodeling in both PT and IT neurons, but that PT neurons are particularly important for mediating its long-term behavioral effects ^14^. In the present study, we used the *Thy1^GFP^* line M mouse, in which GFP expression is largely restricted to infragranular pyramidal neurons ^37^. A limitation of this model is that the genetic identity of the labeled neurons has not been determined. However, previous work suggests that many GFP-positive neocortical neurons in the *Thy1^GFP^* line M mouse are layer 5 PT neurons ^38^.

We focused our studies on the medial frontal cortex, including the ACAd. There are several studies that support a role for ACAd in auditory fear processing. For example, in a trace auditory cued fear conditioning paradigm, ACC neurons were found to have activity related to the termination of fear ^39^. This agrees with some earlier work showing that excitotoxic lesion or transient inactivation of the anterior cingulate cortex impairs tone-shock association ^40^. The anatomical location is important, because ACAd is a relatively large area with anterior and posterior sub-regions. The aforementioned study targeted the anterior portion of the anterior cingulate cortex (dorsal to the prelimbic cortex), which is where we performed imaging. This contrasts with studies that have emphasized anterior cingulate cortex’s role in remote retrieval of contextual fear memory ^41–43^, which tended to target the posterior part (dorsal to the white matter; e.g., ^44^). The anterior portion of ACAd, but to a much lesser extent the posterior portion, sends direct projection to the auditory cortex in mice ^45^. This top-down projection enhances the auditory evoked responses in the auditory cortex in an aversive context ^45^.

Beyond the medial frontal cortex, there is a long history of research implicating the amygdala in fear learning and extinction, including studies examining the role of CREB signaling. Auditory cued fear learning increases signaling in the CREB-CRE transcriptional pathway in the basolateral amygdala ^46^. There is subsequently an elegant series of studies demonstrating that increased CREB expression biases the recruitment of neurons into fear memory ensembles in the lateral amygdala ^47, 48^. However, findings regarding changes in pCREB levels following cued fear extinction have been mixed, with studies reporting different directions of change ^49, 50^.

Causal manipulation studies suggest that pCREB may not be required for the extinction of fear memory ^51^. In the current study, we focused our pCREB analysis on the medial frontal cortex to examine its potential relationship with the changes in dendritic spine turnover observed in this region. However, we want to emphasize that the results do not rule out changes in CREB signaling in response to stress or psilocybin, as the changes may be temporally dynamic, restricted to specific cell populations, or more prominent in other brain regions. Future studies using cell type-specific measurements of CREB activity or broader brain-wide characterization may provide a more comprehensive understanding of the potential responses.

A limitation of the present study is that behavioral testing and longitudinal spine imaging were performed in separate cohorts of mice. In the behavioral cohort, chronic restraint stress impaired fear extinction, consistent with several earlier studies showing that acute or chronic stress blunts the extinction of learned fear associations ^52^. We have body weight as an outcome measure shared between the two cohorts, which suggest the mice experienced similar physiological effects of stress. However, comparable weight changes do not necessarily establish equivalent neurobiological or behavioral effects. Another limitation is that the spine imaging experiment did not include a Control + Psilocybin group. The results allow us to conclude on the effects of psilocybin relative to vehicle following CRS. However, whether chronic stress changes the magnitude or characteristics of the psilocybin-induced structural plasticity remains an important question for future investigation.

Our results align with a couple of earlier studies showing that the rapid acting antidepressant ketamine increases spine density in the context of chronic stress ^8^. Moreover, the non-hallucinogenic 5-HT_2A_ receptor agonist tabernanthalog promotes spine formation in stressed mice ^20^. Relatedly, spine density was elevated in stressed mouse models after repeated administration of LSD ^53^ or single administration of the non-hallucinogenic analog JRT ^22^. The current study adds to this literature by showing that psilocybin, which is currently the most advanced among classic psychedelics in clinical trials, facilitates fear extinction and evokes spine growth in the medial prefrontal cortex in a chronic stress mouse model.

## METHODS

### Animals

For behavioral experiments, we used C57BL/6J mice. For imaging experiments, we used heterozygous *Thy1^GFP^* line M mice (#007788, Jackson Laboratory). A list of all the animals used, including the numbers and experiments, is included in **Table S1**. Animals were 8-12 weeks old at the time of behavioral testing or imaging. Experimental procedures were approved by the Institutional Animal Care & Use Committee at Cornell University (Protocol #2021-0100).

### Chronic restraint stress

Two restraint methods were used: a plastic cone with a nose opening for behavioral experiments, or an acrylic tube for confinement with head fixation for imaging experiments. The latter method was required because the head plate used for two-photon imaging could not fit through the plastic cone. During restraint sessions, mice were individually placed in clean, empty cages without visual contact with other animals for 3-5 hours. Restraints were applied daily for 14 consecutive days (day −14 to day −1). Control mice were handled but not restrained.

### Drugs

Psilocybin was obtained from the Usona Institute’s Investigational Drug and Material Supply Program. The chemical composition of psilocybin was confirmed by high-performance liquid chromatography at Usona Institute. A stock 4 mg/mL solution was prepared by dissolving solid in 0.9% saline, then diluted into a working solution of 0.1 mg/mL. Injections, including vehicle controls, were performed intraperitoneally (i.p.).

### Fear conditions and extinction paradigms

The day after the last restraint session, we began fear learning procedures using a near-infrared video fear conditioning system (MED-VFC2-SCT-M, Med Associates). Prior to each session, mice were brought into a room separated from the fear conditioning system where they habituated for 30 min. Chambers were cleaned between each mouse. On day 0 (conditioning), the chamber had blank straight walls, stainless-steel grid floor, and was cleaned with 70% ethanol (context A). Mice were placed in individual chambers, given 3 min to habituate, then received five presentations of an auditory tone as the conditioned stimulus (CS; 4 kHz, 80 dB, 30 s duration). Each CS co-terminated with a foot shock unconditioned stimulus (US; 0.8 mA, 2 s duration). A 90-s intertrial interval separated the CS-US pairings. On day 2 (extinction), mice were injected with psilocybin (1 mg/kg, i.p.) or saline (10 mL/kg, i.p.) then returned to their home cages for 30 min. They were then placed in conditioning chambers, which were altered with a black plastic A-frame, smooth white floor, and cleaned with cleaned with 1% acetic acid (context B). Each mouse was given 3 min to habituate, then received 15 presentations of the CS without the US separated by a 15-s intertrial interval. On day 10 (renewal), the chamber was altered with a striped curved wall, striped plastic floor, and cleaned with Peroxigard (context C). A subset of mice was placed in the chambers and subjected to the same pattern of CS presentations as day 2. For analysis, freezing was extracted from videos using the VideoFreeze software (motion threshold: 18 au, detection method: linear, minimum freeze duration: 30 frames = 1 s; Med Associates).

### Immunohistochemistry

Mice were sacrificed via transcardiac perfusion immediately following extinction testing on day 2. Tissue samples were fixed with 4% paraformaldehyde, then sectioned at 40 μm using a cryostat (Microm HM550, Thermo Fisher Scientific). Free-floating sections were washed, then blocked for 1 h at room temperature with 5% normal goat serum with 0.3% Triton X-100. Samples were washed, then incubated overnight at 4°C with a 1:1000 dilution of rabbit anti-phospho-CREB (Ser133) (06-519, Sigma-Aldrich) in the blocking solution. Samples were washed and incubated overnight at 4°C with a 1:1000 dilution of conjugated secondary antibody (Alex Fluor 488, goat anti-rabbit; ab150077, Abcam). Sections were washed and mounted on slides with DAPI-containing antifade mounting medium (H-1200-10, Vector labs). Imaging was done at 10X magnification using a wide-field fluorescence microscope (BZ-X810, Keyence). For analysis, a maximum projection image was created from Z-stack images. Cell counting was done using the “Find Maxima” function in ImageJ, then dividing this number by the area of the region analyzed. A minimum of four images were analyzed and averaged for each individual animal. Analysis was performed while blinded to experimental conditions.

### Two-photon imaging

A cranial window was implanted for the *Thy1^GFP^* line M mouse following procedures described previously [5], using 3-mm-diameter glass coverslips and centered at the medial frontal cortex (AP = 1.5 mm, ML = 0.25 mm, relative to bregma). Two-photon imaging experiments were performed using a Movable Objective Microscope (MOM, Sutter Instrument) equipped with a resonant-galvo scanner and a water-immersion objective (XLUMPLFLN, 20x/0.95 NA, Olympus). ScanImage 2020 software was used to control the microscope for image acquisition. To visualize GFP-expressing dendrites, excitation was provided by a femtosecond Axon laser (Axon 920-2 TPC, Coherent). The laser power measured at the objective was <40 mW and a 475-550 nm bandpass filter was used to collect the fluorescence emission. During an imaging session, the mouse was head fixed and anesthetized with 1-1.5% isoflurane. Body temperature was controlled using a heating pad and DC Temperature Controller (40-90-8D, FHC) with rectal thermistor probe feedback. Each imaging session did not exceed 2 hr. We imaged apical tuft dendrites at 0-200 μm below the dura. Multiple fields of view were imaged in the same mouse. For each field of view, 10-40 μm-thick image stackers were collected at 1 μm steps and at 1024×1024 pixels at 0.11 μm per pixel resolution. We kept the same set of imaging parameters for the different imaging sessions. Each mouse was imaged on day −15, −3, −1, 1, 3, 5, and 14. On the day of treatment (day 0), there was no imaging and the mouse was injected with either psilocybin (1 mg/kg, i.p.) or saline (10 mL/kg, i.p.) then returned to its home cage. Analysis was done using ImageJ. Images were processed for motion correction using the StackReg plug-in in ImageJ. If a protrusion extended for >0.4 µm from the dendritic shaft, a dendritic spine was counted. To assess the time course, we calculated the fold change in spine density from baseline (averaging values from sessions on day −3 and day −1) for each dendritic segment. To assess effects of chronic restraint stress, we calculated the fold change in spine density from session on day −15 to baseline for each dendritic segment. To assess changes in spine density due to treatment, we calculated the fold change in spine density from baseline to session on day 14 for each dendritic segment. Dendritic spine scoring was performed while blinded to treatment and time.

### Statistics

Freezing was analyzed using a generalized linear mixed effects model in SPSS (v.29) with fixed factors of stress (control, CRS), drug (vehicle, psilocybin), time (Hab, 1st, 2nd, 3rd, 4th, 5^th^ bin), and their interactions. Individual differences in the overall model were accounted for by including within-subject variation as a random effect term. A diagonal covariance structure was used to account for repeated measures/ Sequential Bonferroni correction was applied to between-group comparisons. The pCREB-positive cell density was analyzed using linear mixed-effects model with stress (control, CRS), drug (vehicle, psilocybin), and brain region (ACAd, PL, ILA) and all their interactions as fixed effects. A random intercept for each mouse was applied to account for repeated regional measurements from the same animal. Models were fit in R (v4.5.1) using the lme4 package (v1.1.37). Fixed effects were tested with Type III F tests using Satterthwaite-approximated degrees of freedom (lmerTest v3.1.3). Analysis of pCREB density was conducted by a trained experimenter who was blinded to treatment conditions. Spine density fold-change across the entire time course was analyzed using a mixed effects model using the lme4 package in R. The model included experimental group (control + vehicle, CRS + vehicle, CRS + psilocybin), time (day −15, −3, −1, 1, 3, 5, 14), sex (male, female), and all second-order interactions as fixed effect terms and a random intercept for dendrites nested by mice. Importantly, variation within mouse and dendrite across days was accounted by including random effects terms for dendrites nested by mice. Post hoc comparisons used estimated marginal means with Tukey correction to contrast psilocybin and saline groups. Spine density fold-change between specific times was tested using a mixed effects model with experimental group (control + vehicle, CRS + vehicle, CRS + psilocybin), sex (male, female), and their second-order interaction as fixed effect terms and a random intercept for dendrites nested by mice.

## ACKNOWLEDGEMENTS

Psilocybin was provided by Usona Institute’s Investigational Drug & Material Supply Program; the Usona Institute IDMSP is supported by Alexander Sherwood, Robert Kargbo, and Kristi Kaylo in Madison, WI. This work was supported in part by a research project sponsored by Intra-Cellular Therapies. The authors acknowledge additional support by NIH/NIMH grants R01MH128217, R01MH137047 (A.C.K.), US Department of Education GAANN Fellowship (A.D.G.), and NIH instrumentation grant S10OD032251 (Cornell Biotechnology Resource Center Imaging Facility).

## AUTHOR CONTRIBUTIONS

C.A.K. and A.C.K planned the study. C.A.K. conducted the imaging experiment and analyzed the data. S.C.W. performed the behavioral and immunohistochemistry experiments, and analyzed the data. A.D.G. assisted with imaging experiment. J.M.S. assisted with immunohistochemistry. C.A.K., S.C.W., and A.C.K. drafted the manuscript. All authors reviewed the manuscript before submission.

